# Growth-inhibiting activity of citronella essential oil to multiple fungal plant pathogens

**DOI:** 10.1101/860718

**Authors:** Mark Angelo O. Balendres, Fe M. Dela Cueva

## Abstract

*Cymbopogon* species are among the most reported essential oils with fungitoxic effect. In this study, mycelial growth of *Fusarium oxysporum* (banana wilt), *Colletotrichum gloeosporioides* (mango anthracnose), *C. falcatum* (sugarcane red rot) and *Neopestalotiopsis* spp. (mango leaf spot) as influenced by varying concentrations of citronella essential oil (CEO) was assessed in *in vitro* assays. Following growth inhibition test, spore germination and germ tube elongation of *C. gloeosporioides* were then examined. Mycelial growth of all test fungal pathogens was strongly inhibited by CEO, but variations were observed among fungal species. This growth inhibition activity was caused by the inhibition of spore germination and germ tube elongation as observed in *C. gloeosporioides*. The findings of this study show the strong growth-inhibitory activity of CEO to multiple fungal pathogens, indicating CEO’s potential as a chemical control approach against these fungal pathogens. Glasshouse and field experiments would establish CEO as one potential alternative to fungicides.

## Introduction

Fungal diseases are among the major contributors to global crop yield losses. The losses are often accompanied by the cost of disease control measures that are usually due to the use of expensive fungicides. Frequent fungicide use can also result in resistance by fungal pathogens (van den Bosch et al., 2011) and may have a negative impact to plants, non-target organisms, and the environment. Consequently, over the last decades, many studies have dealt with alternatives to chemical fungicides (Chitwood, 2002), including but not limited to plant-derived water- (Balendres et al., 2016) and oil-based (Rabari et al., 2018) compounds.

Essential oils and their chemical components are known to have antifungal properties against a wide range of important plant pathogenic species. Among the most notable essential oil-bearing plants are those belonging to the genus *Cymbopogon. Cymbopogon* has 144 species and their oils have been used in cosmetic and pharmaceutical industries (Avoseh et al., 2015). Some *Cymbopogon* species have been also reported to have antifungal properties. Lemongrass oil (e.g. *C. citratus*), for instance, inhibits mycelial growth and spore germination of *Colletotrichum gloeosporioides* from mango (Rabari et al., 2018) and *Fusarium verticillioides* (Sreenivasa et al., 2011). Thus, important *Cymbopogon* species are being grown commercially in tropical and subtropical regions worldwide, including in the Philippines (Camacho et al., 2015), to serve many of its purpose (Avoseh et al., 2015).

Antifungal properties have been reported from six *Cymbopogon* species, namely; *C. citratus, C. martinii, C. nardus, C. flexuosus, C. winterianus* and *C. proximus* (see Table 1). But very little is known of the antifungal activity of *C. winterianus* to plant pathogenic fungi (da Cruz et al., 2015, Dela Cueva & Balendres, 2018). Underscoring the antimicrobial potential of *C. winterianus* against major fungal pathogens in regions where *C. winterianus* is widely grown or commercialized could open up opportunities for a safe, environment-friendly, and cheaper disease control option as compared to fungicides.

**Table 1.** Essential oils of *Cymbopogon* species with *in vitro* antifungal activities to *Colletotrichum gloeosporioides, C. falcatum, Fusarium oxysporum* and *Pestalotiopsis* spp.

| Pathogen | Essential Oil | Some References |
| --- | --- | --- |
| *C. gloeosporioides* | *C. citratus* | Rabari et al (2018)<br>Aquino et al (2012)<br>Cruz et al (2010)<br>Lee et al (2007) |
|  | *C. martinii* | Rabari et al (2018)<br>Rozwalka et al (2010) |
|  | *C. nardus* | Veloso et al (2012) |
|  | *C. flexuosus* | Anaruma et al (2010) |
|  | *C. winterianus* | This study |
| *C. falcatum* | *C. winterianus* | This study |
| *F. oxysporum* | *Species not determined* | Sharma et al (2018) |
|  | *C. citratus* | Sharma et al (2017)<br>Gupta et al (2011)<br>Irkin et al (2019)<br>Lee et al (2007) |
|  | *C. proximus* | Fawzi et al (2009) |
|  | *C. flexuosus* | Pandey et al (2003) |
|  | *C. winterianus* | This study |
| *Pestalotiopsis* spp. | *C. citratus* | Bibiano et al (2017) |
|  | *C. winterianus* | This study |

This study aims to determine the *in vitro* activity of citronella essential oil (*C. winterianus*) to *Fusarium oxysporum* (banana wilt), *Colletotrichum gloeosporioides* (mango fruit anthracnose), *C. falcatum* (sugarcane red rot) and *Neoestalotiopsis* spp. (mango leaf spot), the four fungal pathogens causing diseases of tropical crops.

## Materials and Methods

### Source of Fungal Pathogens

*Fusarium oxysporum* (banana wilt), *Colletotrichum gloeosporioides* (mango fruit anthracnose), *C. falcatum* (sugarcane red rot) and *Neopestalotiopsis* spp. (mango leaf spot) were obtained from the Fungal Repository of the Plant Pathology Laboratory, Institute of Plant Breeding, College of Agriculture and Food Science, University of the Philippines Los Baños (IPB, CAFS, UPLB), Laguna, Philippines. Identity and pathogenicity of these fungal pathogens were previously confirmed by combined traditional (morpho-cultural and pathogenicity characteristics) and molecular (polymerase chain reaction and DNA sequencing) techniques. All fungi were maintained in potato dextrose (PDA) medium (Himedia Laboratories Inc, India).

### Source of Citronella Essential Oil

Pure citronella essential oil (CEO) was purchased from the Land Grant Management Office, UPLB, Laguna, Philippines. The CEO was extracted from *Cymbopogon winterianus* plants grown at the Laguna-Quezon Land Grant, Siniloan, Los Baños Laguna (Camacho et al., 2015), and essential oils were extracted using a steam distillation technique (Pangan et al., 2011).

### *In vitro* Fungal Inhibitory Assay

The effect of CEO on fungal mycelial growth was assessed *in vitro* following the procedure described by (Chen et al., 2014). Citronella essential oil was amended into PDA to achieve the test concentrations 0.25, 1.25 and 2.50 µL/mL PDA. Plates containing PDA without CEO served as a control check. A 5-mm mycelial disc-plug of 6-day old of each fungal pathogen was transferred to CEO-treated and non-treated PDA medium and plates were incubated at 27 °C in 14/10 hours light/dark cycle for 6 days. Radial growth was measured from five-replicate plates. The assay was performed three times.

### Spore Germination and Germ Tube Elongation Assay for *C. gloeosporioides*

Germination and germ tube elongation of *C. gloeosporioides* as influenced by CEO was assessed using harvested conidia, from PDA without CEO, from a 6-day old culture. The mango pathogen was selected as representative fungal species for this assay because it was the most less sensitive fungus in the above assays. The method described by (Chen et al., 2014) was similarly used. One hundred µL of the conidial suspension was spread onto water agar (WA) containing 2.5 µL CEO/mL WA. The test concentration was selected because the growth of all fungal pathogens used in this study was completely inhibited at this rate. Microscopic observations were assessed from 30 randomly selected conidia from four replicate-plates at 0, 2 and 7 days. The spore “germinated” when the length of its germ tube was equal or longer than its length. This assay was performed twice.

### Data Analyses

An analysis of variance (ANOVA) test was performed using the IBM SPSS statistical software (Version 22, IBM, Armonk, NY).

## Results

The growth of all fungal pathogens was significantly inhibited (*P<0.05*) at increasing CEO concentrations (Figure 1). The response of different fungal pathogens to CEO, however, varied depending on the CEO concentrations. The sugarcane pathogen (*C. falcatum*) was the most sensitive with the highest reduction in growth at the lowest concentration tested (0.25 µL), while the mango fruit pathogen (*C. gloeosporioides*) was the most less sensitive. Nevertheless, the growth of all fungal pathogens was suppressed at 2.5 µL CEO.

**Figure 1.**
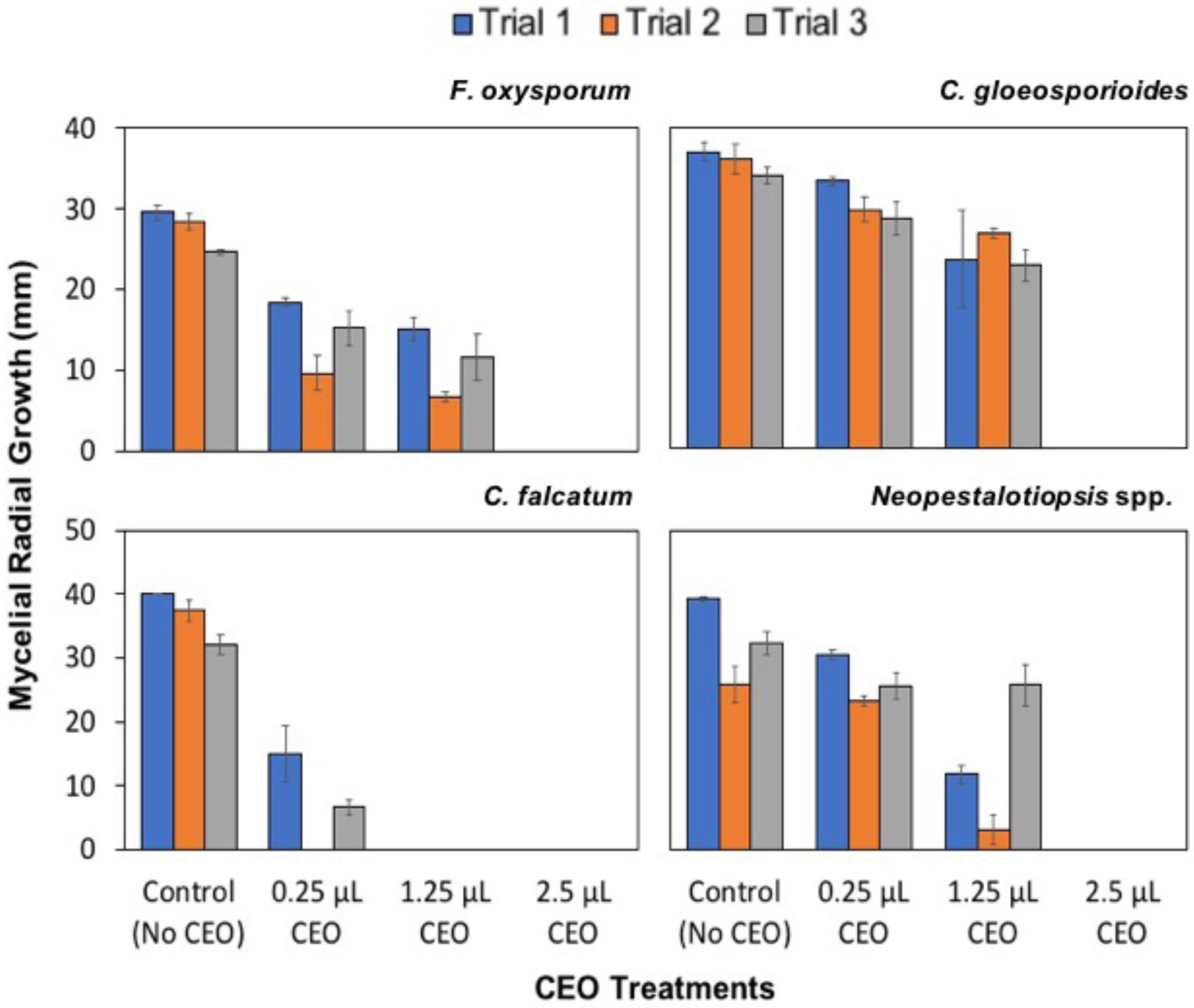
Radial growth (mm) of fungal pathogens *in vitro* after 6 days in various citronella essential oil (CEO) treatments.

When conidia were grown at this rate (2.5 µL CEO), in water agar, germination of *C. gloeosporioides* was inhibited and, subsequently, no germ tube elongated (Figure 2). On the other hand, *C. gloeosporioides* in water agar, without CEO (control treatment), germinated on day 2 and germ tube further elongated on day 7. Additionaly, there was a striking difference between conidia in CEO- and non-CEO treated water ager medium, whereby some conidia in the CEO-treated had size reduced. The conidia in the non-CEO-treated water agar medium swelled and eventually produced germ tubes.

**Figure 2.**
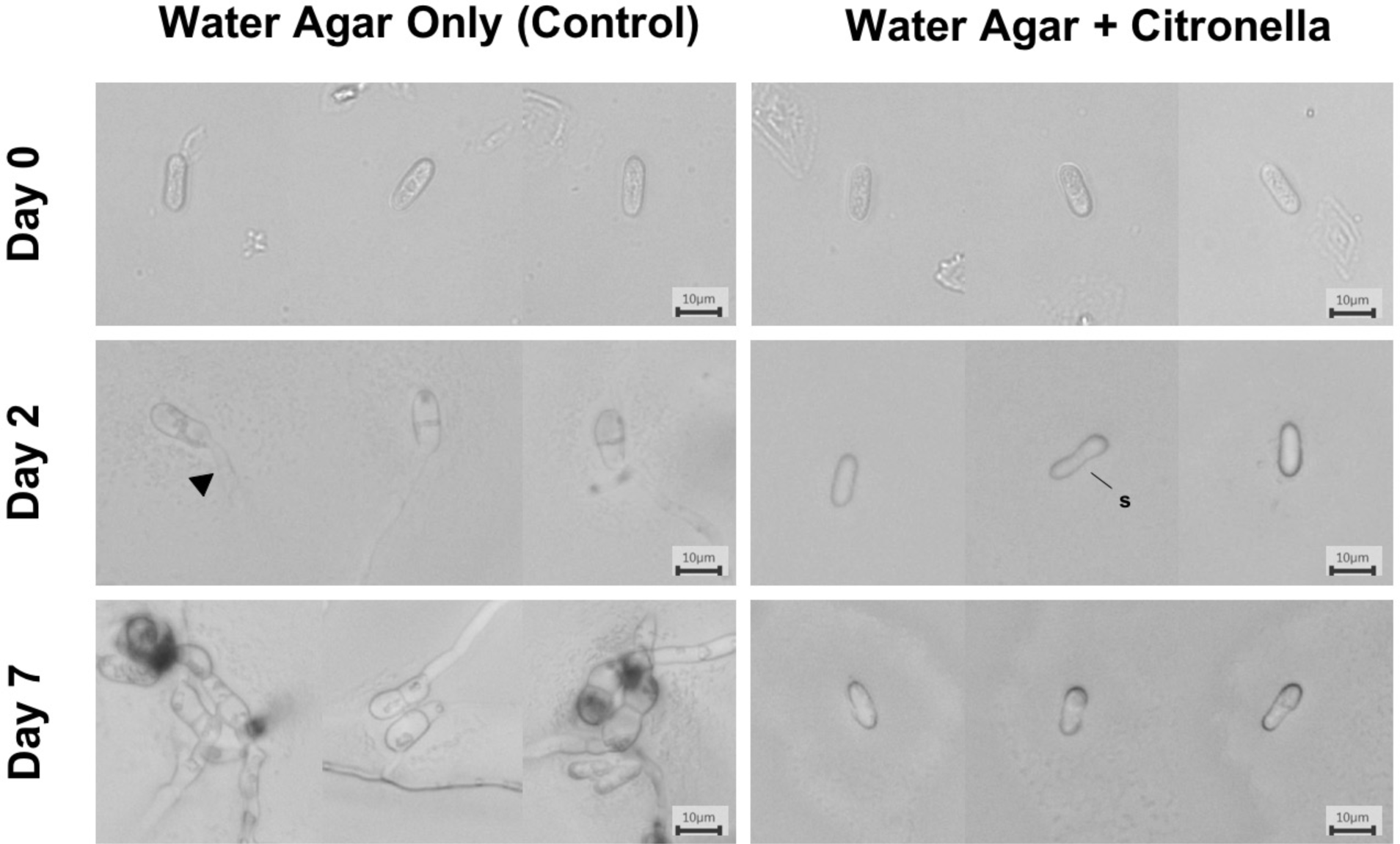
Spore germination and germ tube elongation of *C. gloeosporioides* as influenced by citronella essential oil at 2.5 uL/mL water. Arrowhead, germ tube; s, size reduction.

## Discussion

Essential oils of *Cymbopogon species* have known antifungal properties (see Table 1), but little is known of the activity of citronella (*C. winterianus*) essential oil to other major fungal plant pathogens. This study reports, for the first time, the *in vitro* growth-inhibiting activity of *C. winterianus* to *Fusarium oxysporum* (banana wilt), *C. gloeosporioides* (mango fruit anthracnose), *C. falcatum* (sugarcane red rot) and *Neopestalotiopsis* spp. (mango leaf spot) providing insights into the wide range of pathogens that CEO may be used for in controlling plant diseases. In *C. gloeosporioides*, CEO negatively affects conidia germination and germ tube elongation thus, showing the potent impact of CEO to the biology of the spores, which is an important aspect for *C. gloeosoporioides* reproduction.

da Cruz et al. (2015) and Dela Cueva & Balendres (2018) showed that *C. winterianus* inhibits the growth of *F. solani* (guava decline) and *C. acutatum sensu lato* (chili anthracnose), respectively. This study shows that CEO also inhibits four other fungal pathogens and, more notably, fungal pathogens that cause varying symptoms (fruit anthracnose, leaf spot, red rot, and wilt), suggesting the potential of CEO to control the growth of various fungi irrespective of the fungus pathogenesis. This multiple fungus growth-inhibiting effect is similarly observed on *C. citratus* (lemongrass) and other essential oils (e.g. thyme (Lee et al., 2007)).

Essential oils have direct fungitoxic action to fungal conidia. Rozwalka et al. (2010) observed that conidia of *C. gloeosporioides* and *C. musae* was severely damaged after exposure to lemongrass essential oil and thus, making conidia unable to germinate. In this study, *C. gloeosporioides* conidia indeed did not germinate and did not produce germ tubes after exposure to CEO compared to the control check (no-CEO PDA). Thus, essential oils of *Cymbopogon* species negatively affect the integrity of the cells thereby making conidia unviable.

## Conclusion

Overall, this study demonstrates the growth-inhibiting activity of citronella essential oil (CEO) to *Fusarium oxysporum* (banana wilt), *C. gloeosporioides* (mango fruit anthracnose), *C. falcatum* (sugarcane red rot) and *Neopestalotiopsis* spp. (mango leaf spot) *in vitro*. CEO may be beneficial in suppressing disease development. Further work in the glasshouse and in the field supporting disease-inhibiting and non-phytotoxic activities of CEO could strengthen, harness, and promote CEO as a relatively safe, environment-friendly and cheaper disease control option.

## Acknowledgments

We thank Rizalina Tiongco, Joanne Langres, Fatima Silva, JayVee Mendoza, Meldy Vibal, Cecilia Pascual and Jaime Tumolva for technical assistance.

## Ethical approval

This article does not contain any studies with human participants or animals performed by any of the authors.

## References

Anaruma ND, Schmidt FL, Duarte MCT, et al. (2010) Control of *Colletotrichum gloeosporioides* (penz.) Sacc. in yelow passion fruit using *Cymbopogon citratus* essential oil. Braz J Microbiol 41, 66–73.

Aquino CF, Sales NDP, Soares EPS, Martins ER (2012) Chemical characterization and action of essential oils in the management of anthracnose on passion fruits. Revista Brasileira De Fruticultura 34, 1059–67.

Avoseh O, Oyedeji O, Rungqu P, Nkeh-Chungag B, Oyedeji A (2015) *Cymbopogon* species; ethnopharmacology, phytochemistry and the pharmacological importance. Molecules 20, 7438–53.

Balendres MA, Nichols DS, Tegg RS, Wilson CR (2016) Metabolomes of potato root exudates: compounds that stimulate resting spore germination of the soil-borne pathogen *Spongospora subterranea*. J Agric Food Chem 64, 7466–74.

Bibiano HD, Saber ML (2017) Mycelial growth inhibition of plant pathogenic fungi by extracts. Revista Agrogeoambiental 9, 61–71.

Camacho SH, Carandang AP, Camacho LD, et al. (2015) Economic potential of small-scale citronella (*Cymbopogon winterianus*) production in the Philippines. Phil J Crop Sci 40, 73–81.

Chen F, Han P, Liu P, Si N, Liu J, Liu X (2014) Activity of the novel fungicide SYP-Z048 against plant pathogens. Sci Rep 4.

Chitwood D (2002) Phytochemical based strategies for nematode control. Annu Rev Phytopathol 40, 221–49.

Cruz MJD, Clemente E, Cruz MED, Mora F, Cossaro L, Pelisson N (2010) Effects of bioactive natural compounds on the postharvest conservation of mango fruits cv. Tommy Atkins. Ciencia E Agrotecnologia 34, 428–33.

Da Cruz TP, Alves FR, Mendonca RF, et al. (2015) Fungicide activity of essential oil *Cymbopogon winterianus* Jowit (citronella) against *Fusarium solani*. Biosci J 31, 1–8.

Dela Cueva F, Balendres MA (2018) Efficacy of citronella essential oil for the management of chilli anthracnose. Eur J Plant Pathol 152, 461–468.

Fawzi EM, Khalil AA, Afifi AF (2009) Antifungal effect of some plant extracts on *Alternaria alternata* and *Fusarium oxysporum*. Afr J Biotech 8, 2590–7.

Gupta A, Sharma S, Naik SN (2011) Biopesticidal value of selected essential oils against pathogenic fungus, termites, and nematodes. Int Biodet Biodeg 65, 703–7.

Irkin R, Korukluoglu M (2009) Effectiveness of *Cymbopogon citratus* L. essential oil to inhibit the growth of some filamentous fungi and yeasts. J Medicinal Food 12, 193–7.

Lee SO, Choi GJ, Jang KS, Lim HK, Cho KY, Kim JC (2007) Antifungal activity of five plant essential oils as fumigant against postharvest and soilborne plant pathogenic fungi. Plant Pathol J 23, 97–102.

Pangan R, Rodulfo V, Gallegos R, Amongo R, Arenal A (2011) Improving the operation of the conventional citronella (*Cymbopogon nardus*) oil extractor through optimization studies. Agric Mech J 18, 1.

Rabari VP, Chudashama KS, Thaker VS (2018) *In vitro* screening of 75 essential oils against *Colletotrichum gloeosporioides*: a causal agent of anthracnose disease of mango. Int J Fruit Sci 18, 1–13.

Rozwalka LC, Alves E, Do Amaral DC (2010) Ultrastructural study of conidia of *Colletotrichum gloeosporioides* and *Colletotrichum musae* treated with essential oils. Interciencia 35, 912–5.

Sharma A, Rajendran S, Srivastava A, Sharma S, Kundu B (2017) Antifungal activities of selected essential oils against *Fusarium oxysporum* f. sp. lycopersici 1322, with emphasis on *Syzygium aromaticum* essential oil. J Bioscience Bioengin 123, 308–13.

Sharma A, Sharma NK, Srivastava A, et al. (2018) Clove and lemongrass oil based non-ionic nanoemulsion for suppressing the growth of plant pathogenic *Fusarium oxysponum* f.sp *lycopersici*. Indust Crops Prod 123, 353–62.

Sreenivasa MY, Dass RS, Raj APC, Prasad MNN, Achar PN, Janardhana GR (2011) Assessment of the growth inhibiting effect of some plant essential oils on different Fusarium species isolated from sorghum and maize grains. J Plant Dis Prot 118, 208–13.

Van Den Bosch F, Paveley N, Shaw M, Hobbelen P, Oliver R, 2011. The dose rate debate: does the risk of fungicide resistance increase or decrease with dose? Plant Pathology 60, 597–606.

Veloso RA, De Castro HG, Cardoso DP, Dos Santos GR, Barbosa LCD, Da Silva KP (2012) Composition and fungitoxicity of essential oil of citronella grass as affected by organic fertilization. Pesquisa Agropecuaria Brasileira 47, 1707–13.

